# Impact of specimen age on its DNA quality for Formalin-Fixed Paraffin-Embedded HPV specimens

**DOI:** 10.1101/420224

**Authors:** James Yeongjun Park

## Abstract

Formalin fixation and paraffin embedding (FFPE) allows the storage of diagnostic and surplus tissue in archival banks. Therefore, FFPE is now a standard method for long-term preservation of tissue biopsies as FFPE of samples preserves the morphology of tissue. Unfortunately, the FFPE process engenders chemical changes and degradation in tissue macromolecules that can pose threats to reliable subsequent analysis. DNA, while more resistant to FFPE in comparison to RNA and protein, is also subject to such chemical formations and degradation. This study provides robust findings about the relationship between DNA quality and specimen age from 10252 FFPE HPV specimens. This paper suggests that there is a perceptible degrading effect in DNA quality as specimens age. This study suggests that the biospecimen may begin to take on new characteristics after certain storage years, and such changes may result in inaccurate determinations of the molecular characteristics of the biospecimen during analysis. Results from this study demonstrate that older HPV specimens are more inclined to be tested negative or inadequate from commercial HPV genotyping assays. Same findings are conferred when multiple genotyping assays are involved with HPV testing. Also, older HPV samples typify fewer HPV types compared to younger HPV samples. The results from this study will be useful to enhance potential scope for next fixation methods such as ethanol fixation that may be equally useful for both molecular profiling and histology as FFPE.

## Introduction

With the sizable expansion in the human population, there will be a large increase in biospecimen. There has been an extraordinary evolution of next-generation technologies that allow targeted evaluation of biospecimen, and the availability of diverse human biospecimen to aid biomedical research will continue to be critical to advancing the state of knowledge regarding the origins and progression of diseases such as cancer. With this advancement in mind, provision of tissue necessitates a systematic approach to ensure that the tissue does not degrade to a point where the specimen’s viability is damaged. Although these specimens have yielded countless insights and advanced the status of disease pathogenesis, there is increasing scientific need for large biorepository and appropriate quality controls of such specimens.

The disadvantage of conducting biomarker profiling in fresh-frozen tissue taken at the time of surgery is that the tissue might not be entirely representative of tumor in addition to the lack of adequate numbers and follow-up to make clear conclusions as to a biomarker’s prognostic potential. In contrast, Formalin fixation and paraffin embedding (FFPE) allows the storage of diagnostic and surplus tissue in archival banks. Therefore, FFPE is now a standard method for long-term preservation of tissue biopsies as FFPE of samples preserves the morphology of tissue, and many FFPE samples are archived in biorepository. The aim of most biorepositories is to collect, process, store, and distribute human biospecimen for use in basic, translational, and clinical research, which requires the highest standards of operation as modern research heavily relies on high-quality human biospecimen. DNA isolated from FFPE can be used for amplification of short amplicons, which is appropriate for the downstream analysis by the polymerase chain reaction (PCR) in combination with oligonucleotide probe procedures. Unfortunately, the FFPE process engenders chemical changes and degradation in tissue macromolecules that can pose threats to reliable subsequent analysis. DNA, while more resistant to FFPE in comparison to RNA and protein, is also subject to such chemical formations and degradation. Few studies have corroborated that not all preservation or fixation methods render DNA that is suitable for subsequent amplification [9].

Due to advancements in understanding how to best cryopreserve human tissues, samples can now be effectively cryopreserved for years and still be viable specimens for studying many cellular process; however, not many studies have yet investigated exactly how long these samples stay high quality. One study assessed the effects of fixation on subsequent DNA amplification to test the effects of specimen age. The results from this study showed that after 16 years, 90 percent of samples were suitable for the amplification of 268-bp B-globin fragment. Successful amplification of this fragment decreased to 45 percent for 20-year-old samples [9]. This study provides more robust findings about the relationship between DNA quality and specimen age from a much larger number of samples; it suggests that there is a perceptible degrading effect in DNA quality as specimens age. Our findings suggest that the biospecimen may begin to take on new characteristics after certain storage years, and such changes may result in inaccurate determinations of the molecular characteristics of the biospecimen during analysis.

## Method

Of 13,144 HPV specimens, analyses are performed on 10,252 samples that have their epidemiology data available. Specimen age of samples is measured by the amount of time elapsed between samples’ initial receipt date and their extraction date. Three genotyping assays are used to typify HPV types of these samples. A 10 µl aliquot of the purified DNA is first tested with the linear array (LA). LA detects 37 high and other-risk genotypes individually. Although this assay is useful in detecting many HPV types with genotype-specific probes, the diversity of the HPV spectrum makes it difficult to precisely identify individual genotypes. Therefore, samples with HPV negative or inadequate LA results are re-tested with the INNO-LiPA and the LBP LiPA assays as their short PCR target lengths may be more adequate for low quality DNA from FFPE tissues. From the Linear Array, 81.1% of the samples tested positive (n = 10660) while 10.1% tested negative (n = 1327) and 8.8% inadequate (n = 1157). When the LiPA assay is incorporated. 96.4% of the samples tested positive (n = 12670) while 3.1% tested negative (n = 395) and 0.6% tested inadequate (n = 79). This algorithm of testing and reporting of results of three commercial assays for HPV detection and typing in FFPE samples is shown below.

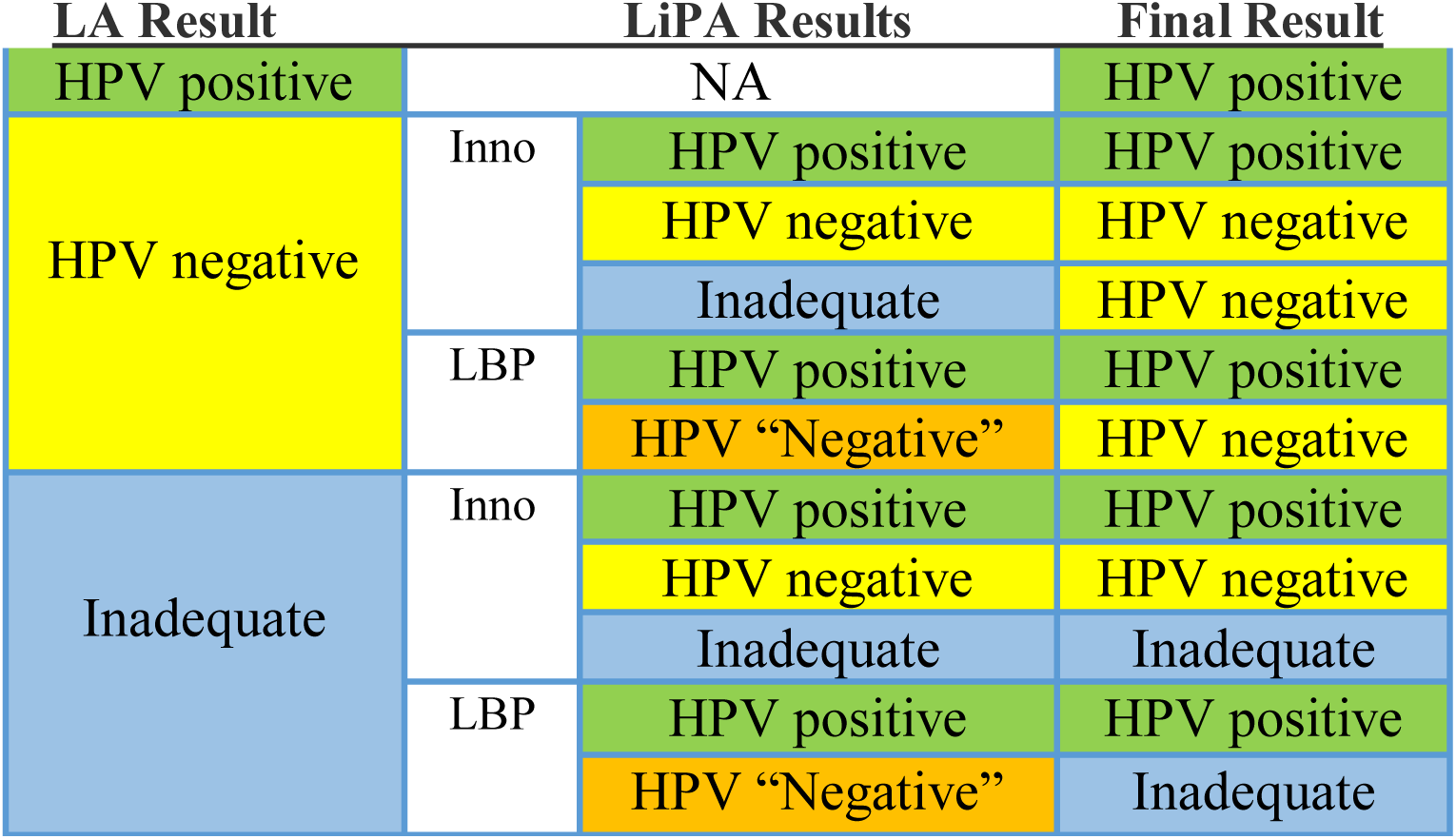

| LA Result |  | LiPA Results | Final Result |
| --- | --- | --- | --- |
| HPV positive |  | NA | HPV positive |
| HPV negative | Inno | HPV positive | HPV positive |
|  |  | HPV negative | HPV negative |
|  |  | Inadequate | HPV negative |
|  | LBP | HPV positive | HPV positive |
|  |  | HPV "Negative" | HPV negative |
| Inadequate | Inno | HPV positive | HPV positive |
|  |  | HPV negative | HPV negative |
|  |  | Inadequate | Inadequate |
|  | LBP | HPV positive | HPV positive |
|  |  | HPV "Negative" | Inadequate |

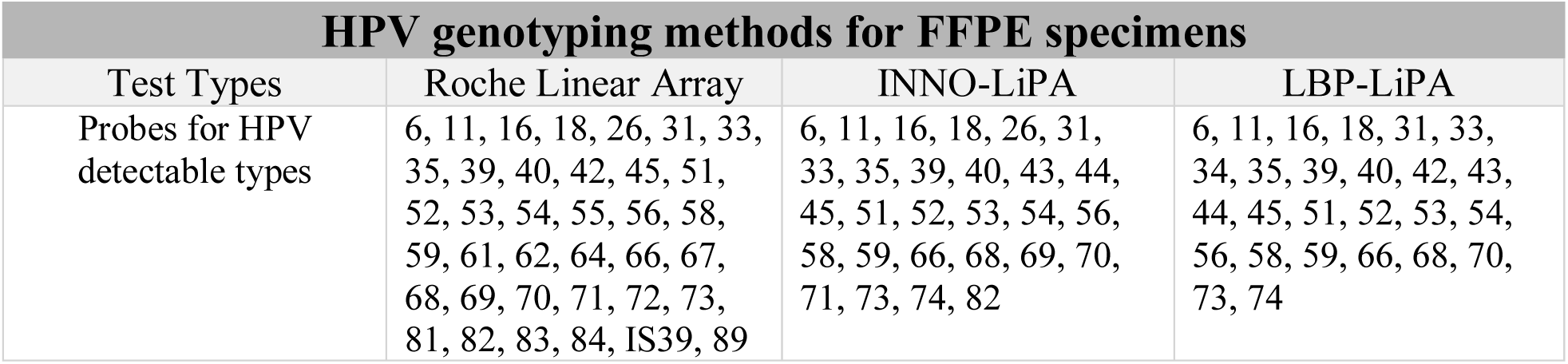

## Result

Analyses are performed in two test sets. The first analysis investigates assay proportion of HPV result for 10,252 samples from the LA assay alone while the second analysis combined the LiPA assay in conjunction with the LA assay. Multinomial logistic regression is used for this analysis as assay status typifies three possible discrete outcomes. This model predicts the possibilities of the different outcomes of a categorically distributed dependent variable given an independent variable of specimen age measured in year.

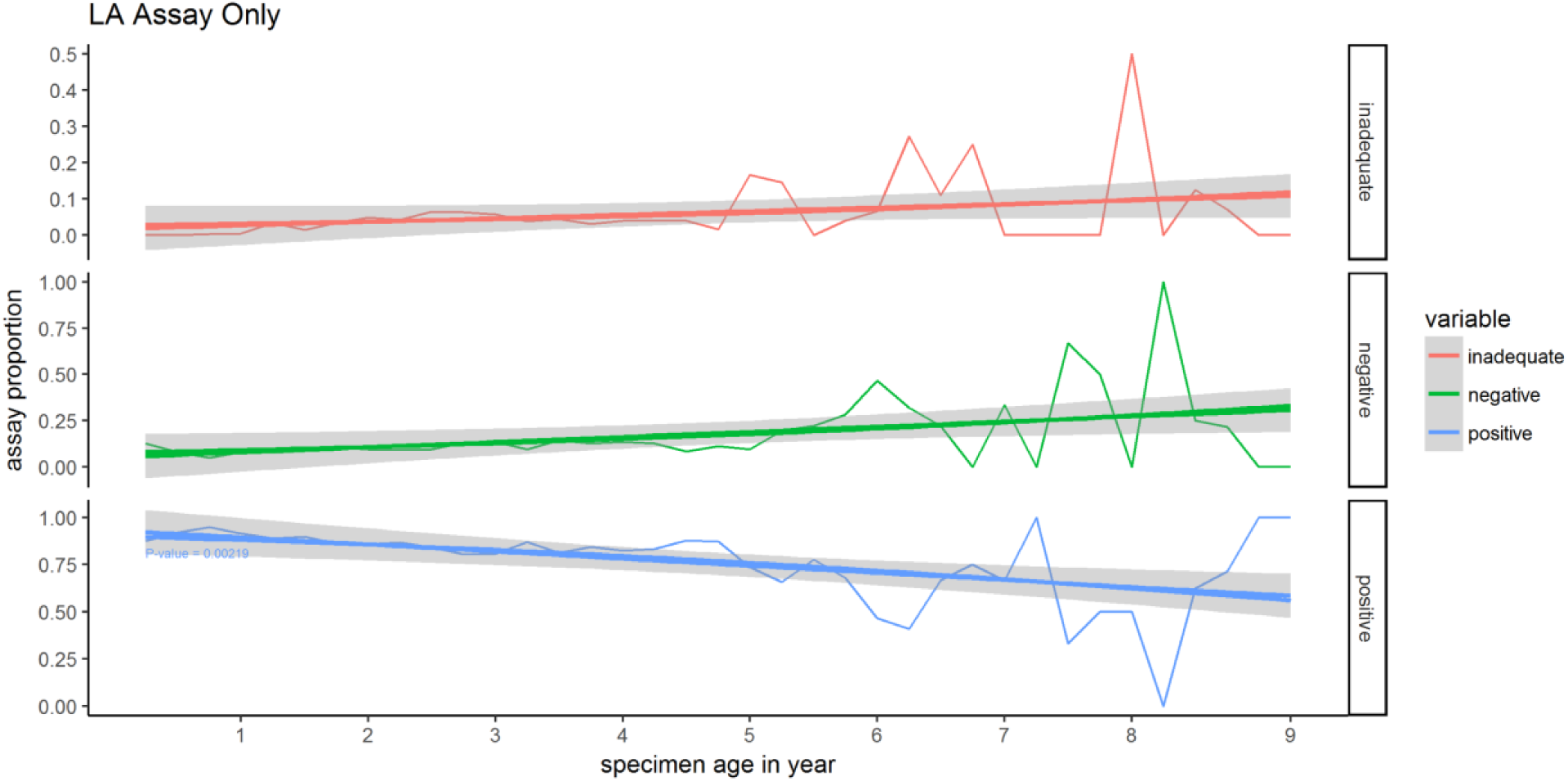

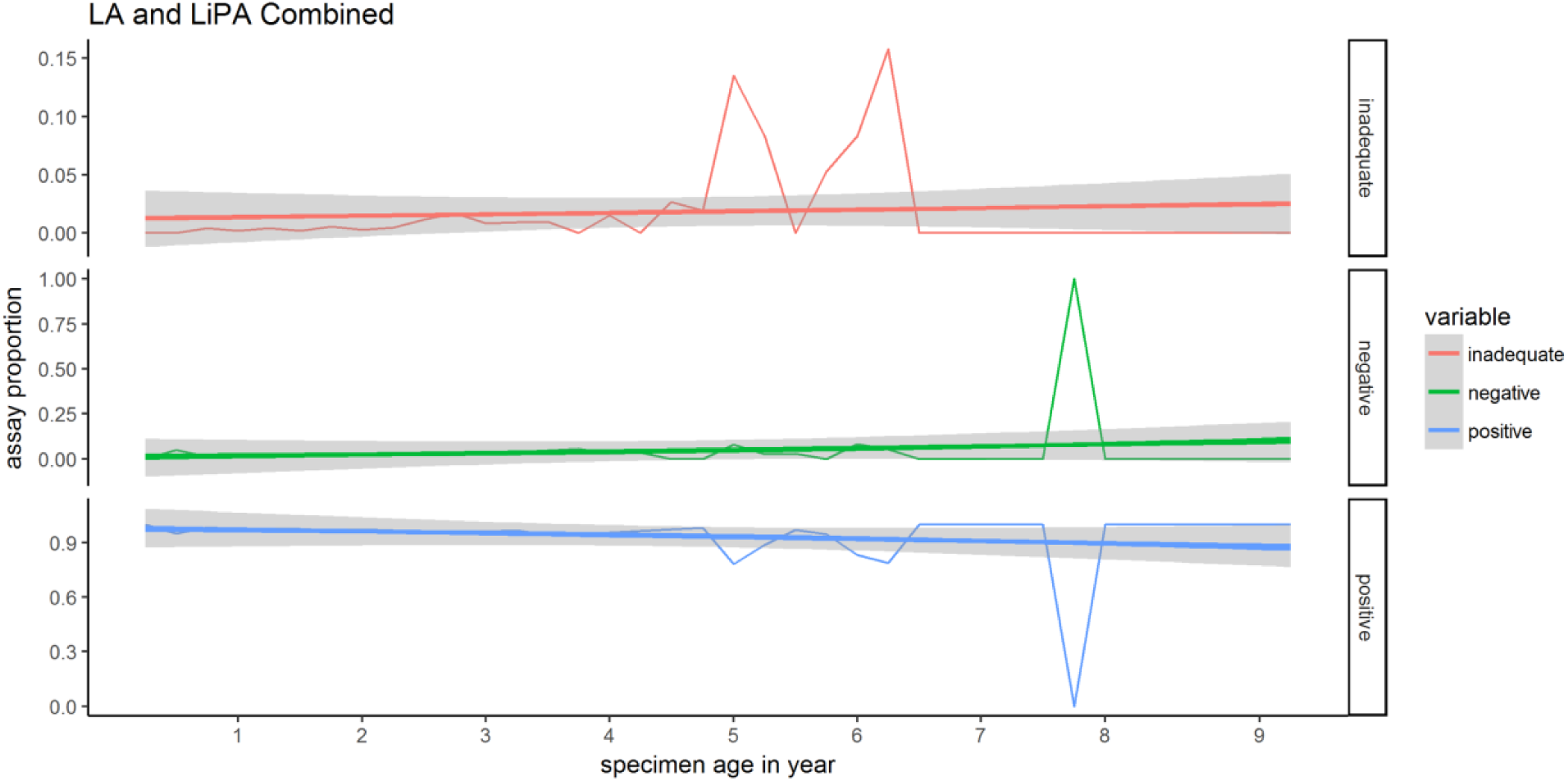

From both test sets, positive coefficients are obtained for HPV negative and inadequate samples while stronger negative coefficients are found for HPV positive trends. Most samples’ age ranges from one to five, and very few samples were seven years old or older proportionally, which explains the seeming spikes in the second figure for samples that are seven to eight years old.

Another analysis is performed to examine a potential relationship between the total number of HPV types detected per sample and specimen age. Samples that are five years or older are categorized as “5+” as there are proportionally much fewer samples that are five years or older. Note that the LA assay detects 37 HPV types while the LiPA assay detects fewer HPV types. The LiPA assay also detects several HPV types that are not detectable by the LA assay so the total number of HPV type detected is increased when the LiPA assay is examined in addition to the LA assay. Both negative and inadequate samples are classified as samples with zero type.

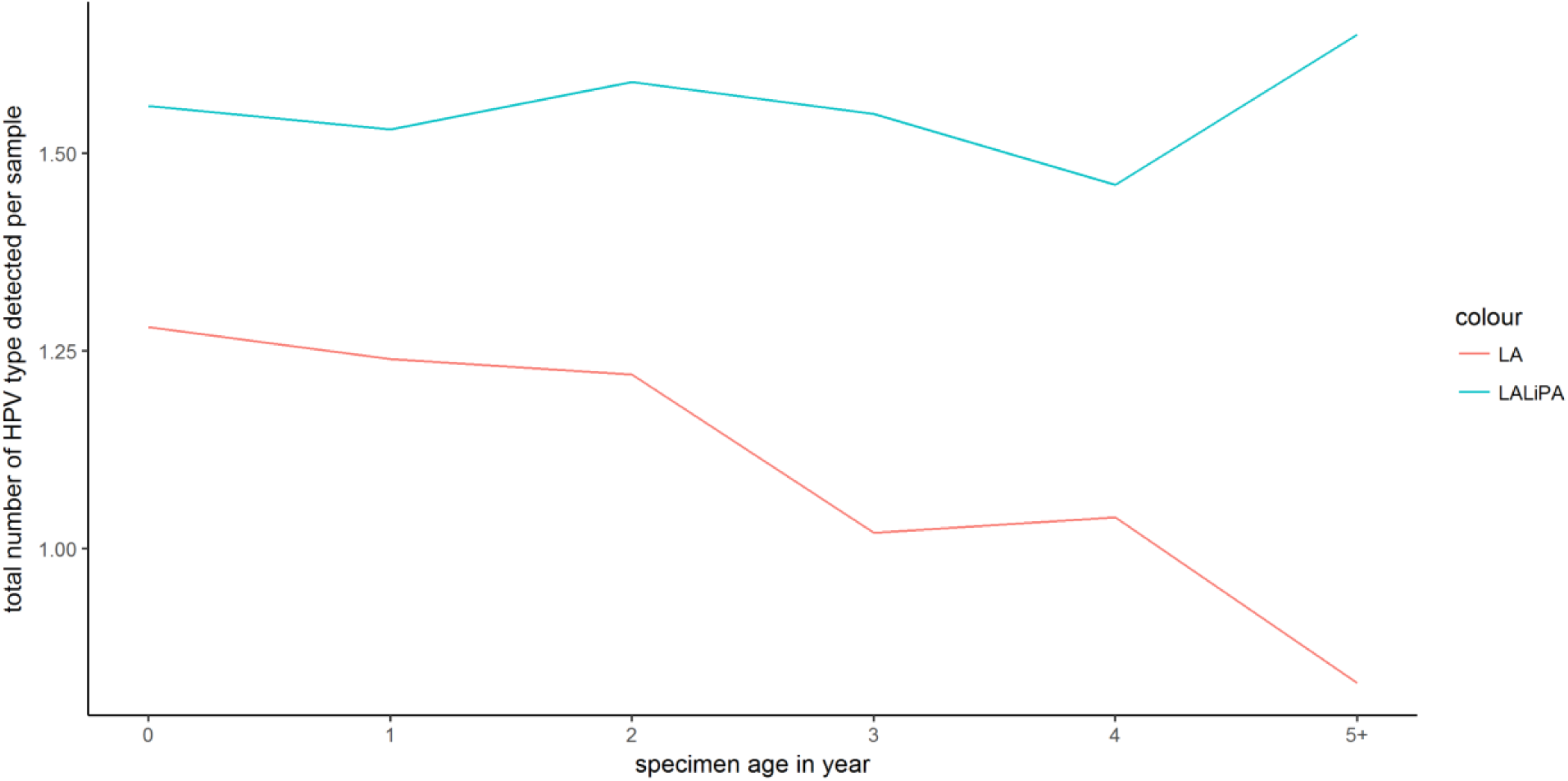

Striking results are obtained when only the LA assay is examined; as samples get older, they linearly typify fewer HPV types. The results are less drastic when the LiPA assay is incorporated in conjunction with the LA assay; this may be due to having fewer samples that are four years or older in comparison to samples that are up to four years old. These results evince that FFPE samples may degrade after certain storage years, which may lead to inaccurate determinations of the assay results.

## Conclusion

The aim of any biospecimen sample should be to maintain and disseminate the highest quality on the intended research use in which the high-quality specimen stimulates the biology of the specimen prior to its removal from the human participant. Our findings suggest that the biospecimen may begin to take on new characteristics after certain storage years, and such changes may result in inaccurate determinations of the molecular characteristics of the biospecimen during analysis. DNA is deemed the most resistant to FFPE degradation of all macromolecules, but our results indicate that DNA may undergo significant degradation as samples age. Ongoing refinements in laboratory technique and tissue fixation are likely to improve in furtherance. The results from this study will be useful to enhance potential scope for next fixation methods such as ethanol fixation that may be equally useful for both molecular profiling and histology as FFPE. Over many years, a multitude of guidelines have been generated based on operations and insights to standardize biorepository operations. More studies are needed to construct an optional fixation standard operating procedure so that the variability in degradation is maximally reduced.

